# A RNA-seq characterization of the porcine sperm microbiome

**DOI:** 10.1101/2020.03.16.994244

**Authors:** M. Gòdia, Y. Ramayo-Caldas, L. M. Zingaretti, S. López, J. E. Rodriguez-Gil, M. Yeste, A. Sánchez, A. Clop

## Abstract

The microbiome plays a key role in homeostasis and health and it has been also linked to fertility and semen quality in several animal species including swine. Despite the more than likely importance of sperm bacteria on the boar’s reproductive ability and the dissemination of pathogens and antimicrobial resistance genes, a high throughput characterization of the swine sperm microbiome remains undone. The current study aimed at profiling the boar sperm bacterial population and its relationship with seven semen quality traits.

We carried RNA-seq on 40 ejaculates and we found that it contains a broad population of bacteria. The most abundant phyla were *Proteobacteria* (39.1%), *Firmicutes* (27.5%), *Actinobacteria* (14.9%) and *Bacteroidetes* (5.7%). The predominant species contaminated sperm after ejaculation from soil, faeces and water sources (*Bacillus megaterium*, *Brachybacterium faecium*, *Bacillus coagulans*). Some potential pathogens were also found but at relatively low levels (*Escherichia coli*, *Clostridioides difficile*, *Clostridium perfringens*, *Clostridium botulinum* and *Mycobacterium tuberculosis*). We also identified 3 potential antibiotic resistant genes from *E. coli* against chloramphenicol, *Neisseria meningitidis* against spectinomycin and *Staphylococcus aureus* against linezolid. None of these genes were highly abundant. Finally, we classified the ejaculates into categories according to their bacterial features and semen quality parameters and identified two categories that significantly differed for 5 semen quality traits and 13 bacterial features including the genera *Acinetobacter*, *Stenotrophomonas* and *Rhodobacter*. Our results show that boar semen contains a rich microbiome with potential pathogens and antibiotic resistance genes which may affect its reproductive performance.

## Introduction

Scientific research has lead to the discovery that microbiota, the microorganisms present in the different compartments in an animal organism, play a crucial role in physiological homeostasis and health (Schroeder and Bäckhed, 2016, Mahboubi et al., 2016, Moreno et al., 2016) including sperm quality and male fertility (Weng et al., 2014, Baud et al., 2019). The male’s reproductive ability is represented by a set of traits that are important for human health and for the efficiency and sustainability of animal production. In swine, semen quality is regularly measured in the artificial insemination studs as a proxy of the fertilization ability of that sample. Growing research is being devoted to understanding the biological basis and identifying molecular markers linked to semen quality in humans and other animal species. As the presence of bacterial communities in ejaculates is common and the microbiome is popping up as a big contributor of a broad range of phenotypes, several studies have been carried in the field of men fertility (Mändar et al., 2015, Weng et al., 2014, Hou et al., 2013) and boar sperm quality (Úbeda et al., 2013, Sepúlveda et al., 2016). Weng et al. (Weng et al., 2014) identified a complex population of bacteria in human sperm but most interestingly, found that the abundance of some bacteria was related to male fertility. *Lactobacillus crispatus*, *Gardnerella vaginalis* and *Lactobacillus acidophilus* were more abundant in the fertile samples whilst *Prevotella vibia* and *Haemophilus parainfluenzae* were present at higher proportion in the unfertile sperm (Weng et al., 2014). In a more recent study, a group led by Stephen Krawetz (Swanson et al., 2020) used sperm RNA-seq datasets to identify transcripts of bacterial origin and shed light to the bacterial composition of an ejaculate. They found a diverse bacterial population mostly characterized by members of the phyla *Firmicutes*, *Proteobacteria*, *Bacteroidetes* and *Actinobacteria* (Swanson et al., 2020).

In pigs, the presence of bacteria in sperm is well documented and bacterial populations in ejaculates are common (Althouse and Lu, 2005). In pigs, most of the bacteria present in semen ejaculates seem to have an external origin and might thus have contaminated the sperm after ejaculation (Schulze et al., 2015). Ubeda and co-authors, using cell culture, concluded that the most abundant bacteria in pig semen were from the Enterobacteriaceae family and included, in order of abundance, *Serratia marcescens*, *Klebsiella oxytoca*, *Providencia stuartii*, *Morganella morganii*, *Proteus mirabilis*, and *Escherichia coli*. Among these, *S. marcescens*, *K. oxytoca*, *M. morganii*, or *P. mirabilis* were, in an individual basis, negatively associated with sperm quality (Úbeda et al., 2013). Schulze also recently identified the presence of several species of *Lactobacillus* and an association, *in vitro*, between the abundance of *Lactobacillus buchneri* with sperm motility, mitochondrial activity and membrane integrity and *Lactobacillus animalis* with motility (Schulze et al., 2018). To control bacterial growth in sperm, antimicrobial agents are commonly added as essential components to semen extenders (Althouse and Lu, 2005). Nonetheless, bacteria in these extended ejaculates can be still present due to incomplete efficiency of the antibiotics which could be partially caused by the expression of antimicrobial resistance genes (ARG) by these bacteria. Current high throughput sequencing technologies provide unprecedented capacity to study microbial ecosystems. However, it has not yet been applied to characterize the pig sperm microbiome. In the current study, we characterized the boar sperm microbiome by RNA-seq on ejaculates of 40 pigs and indicate the existence of a potential link between the sperm microbiome and semen quality traits.

## Materials and methods

### Sample collection

Specialized professionals obtained fresh ejaculates from 40 Pietrain boars from three different commercial farms, with the hand glove method. Ejaculates were collected between March 2015 and January 2017 and boar ages ranged from 9 to 55 months old. The fresh ejaculates were preserved in Androstar^®^ Plus boar semen extender that contains the following antibiotics: apramycin sulphate, cephalosporin – third generation - and gentamicin sulphate. Sperm ejaculates were phenotyped as in (Gòdia et al., 2018). Then, they were purified using the BoviPure^™^ colloidal silica particles reagent (Nidacon; Mölndal, Sweden) as detailed by Gòdia et al. to obtain only mature spermatozoa cells (Gòdia et al., 2018).

### RNA extraction, qPCR validation, library prep, sequencing

RNA from spermatozoa pellets was extracted using a standard Trizol^®^ approach and treated with TURBO DNA-free™ Kit (Invitrogen; Carlsbad, USA) (Gòdia et al., 2018). RNA samples were subjected to RT-qPCR assays to validate the absence of RNA from contaminating diploid cells (mainly leukocytes and keratinocytes) and genomic DNA (Gòdia et al., 2018). Total RNA was subjected to ribosomal RNA (rRNA) depletion with the Ribo-Zero Gold rRNA Removal Kit (Illumina, CA, USA). RNA-seq libraries were prepared with SMARTer Universal Low Input RNA library Prep kit (Clontech, France) and sequenced in an Illumina’s HiSeq2000/2500 system to generate 75 base pair long paired end reads. Ten of these RNA-seq datasets have been analysed to characterize the boar sperm transcriptome in our previous study (Gòdia et al., 2019). The RNA-seq data used in this study (total RNA-seq runs) is accessible at the NCBI’s SRA within the under the SRA study accession SRP183646.

### Identification of RNA molecules of bacterial origin

RNA-seq reads of low quality and adaptor contaminations were removed with Trimmomatic v.0.36 (Bolger et al., 2014). Filtered reads were then mapped to the *Sus scrofa* genome (Sscrofa11.1) with HISAT2 v.2.1.0 (Kim et al., 2015) with default parameters except “--max seeds 30” and “-k 2”. The reads that did not map to Sscrofa11.1 were screened against the catalogue of porcine Transposable Elements from the Repbase database (Bao et al., 2015) with HISAT2 v.2.1.0 (Kim et al., 2015). The reads that remained unmapped were taxonomically classified and quantified with Kraken v.0.10.5 (Wood and Salzberg, 2014) with a threshold score of 0.15 and using the default database that includes NCBI taxonomic information and complete genomes from RefSeq of archaeal, bacteria, phage and viral domains. Only the bacterial-assigned reads were kept for further analysis. The number of reads assigned to a given taxon was normalized by sequencing depth, as counts per million (CPM). The list of potential pathogens in swine was extracted from the Professional Pig Community pig333 site (www.pig333.com/pig-diseases). This site includes 24 bacterial-borne diseases.

### Detection of antimicrobial resistance genes

Unmapped reads were also subjected to identification and relative abundance quantification of ARGs. ARGs were identified using BLASTN v.2.7.1 (Camacho et al., 2009) with 100% percentage identity using the Comprehensive Antibiotic Resistance Database (CARD) v.3.0.0 (Jia et al., 2017). The number of reads for each ARG was normalized by sequencing depth, as CPM. The read coverage across ARGs of point mutations was individually visualized using R v.3.5.3 (R Core Team, 2017).

### Relation between bacterial abundance and semen quality traits

Seven sperm phenotypes were measured in the 40 samples as previously described (Gòdia et al., 2018). These included the percentage of viable sperm cells after 5 minutes of incubation at 37 °C (VIAB_5), the percentage of viable sperm cells after 90 min incubation at 37 °C (VIAB_90), percentage of cells with abnormal acrosomes after the 5 min (ACRO_5) and the 90 min (ACRO_90) incubation, the percentage of motile cells measured with the computer-assisted semen analysis (CASA) system (Integrated Sperm Analysis System V1.0; Proiser) after 5 min (MT_5) and 90 min (MT_90) incubation and the percentage of viable cells after an osmotic stress (ORT, Osmotic Resistance Test). All these phenotypes were corrected using the R function “lm” (R Core Team, 2017) for the fixed variables: farm, age and season/year. To calculate the relationship between the semen quality traits and the bacterial features of the sperm microbiome we used Link-HD (Zingaretti et al., 2019), a tool for the integration of heterogeneous data that considers the compositional nature of microbiome datasets. This analysis included the 7 corrected phenotypes and the bacterial features with average CPM ≥ 1 and representing more than 0.001% of the total bacterial read counts.

The phenotypic values of the clusters obtained by Link-he abundances of the 733 bacterial features of the cluster compromises obtained by Link-HD were then compared using the fitZig function of the metagenomeSeq package v. 1.28.2 (Paulson et al., 2013). This function uses a Zero-Inflated Gaussian (Zig) distribution to model the log2-abundances of the Operational Taxonomic Units OTUs) and it is calculated through an Expectation Maximisation (EM) algorithm.

## Results

### RNA-seq statistics

We carried RNA-seq on 40 ejaculates each from a different Pietrain pig and obtained an average of 40.7 M reads per sample. In average, 98.5% of the reads passed the quality control and 82.7% mapped to the porcine genome (Sscrofa11.1). A tiny proportion (0.012%) of the unmapped reads aligned to Repbase (Bao et al., 2015) and 25.1% (an average of 1.7 M reads per library) mapped to microbial genomes with Kraken (Table S1).

### Description of the boar sperm microbiome

We identified 733 bacterial features with average abundance ≥ 1 CPM and representing more than 0.001% of the total bacterial read counts. The total bacterial abundance across samples varied between 2,241 and 180,624 CPMs (Fig. 1 and Table S2). The average and median abundances of bacterial reads were 20,149 and 9,785 CPM, respectively and 3 ejaculates had more than 70,000 bacterial CPM (Fig. 1). The bacterial features included 15 phyla (Fig. 2 and Table S2). The most abundant phyla were *Proteobacteria*, with an average of 39.1% of bacterial reads, *Firmicutes* (27.5%), *Actinobacteria* (14.9%) and *Bacteroidetes* (5.7%) (Fig. 2 and Table S2). At the species level, the analysis identified 254 bacterial species (Table S2). The most abundant species were, in this order, *Bacillus megaterium* (868 CPMs and 4.3% of the bacterial reads), *Brachybacterium faecium* (3.3%), *Bacillus coagulans* (1.2%) and *Campylobacter hominis* (1.0%) (Table 1).

**Figure 1.**
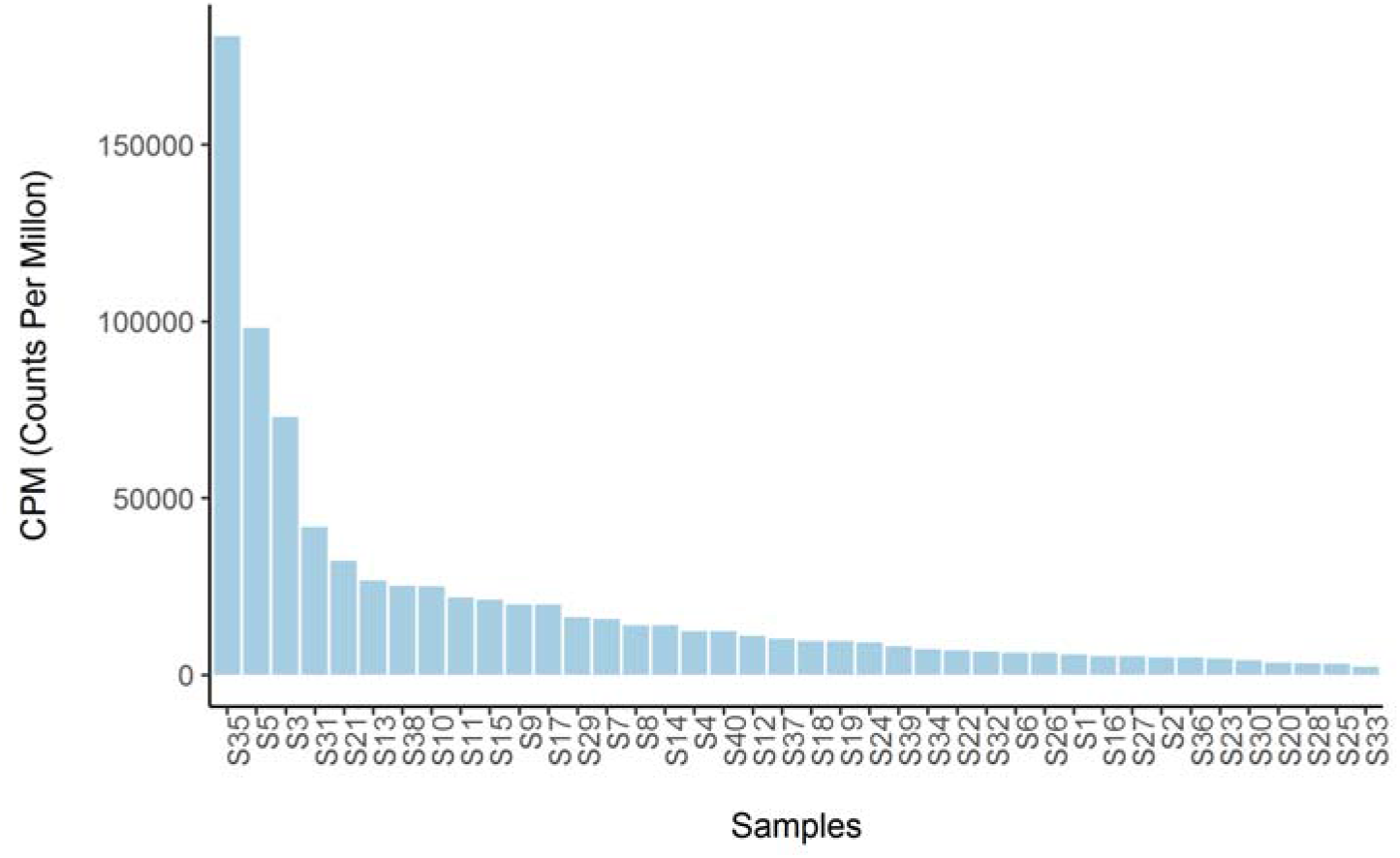
Distribution of overall bacteria abundance for each animal.

**Figure 2.**
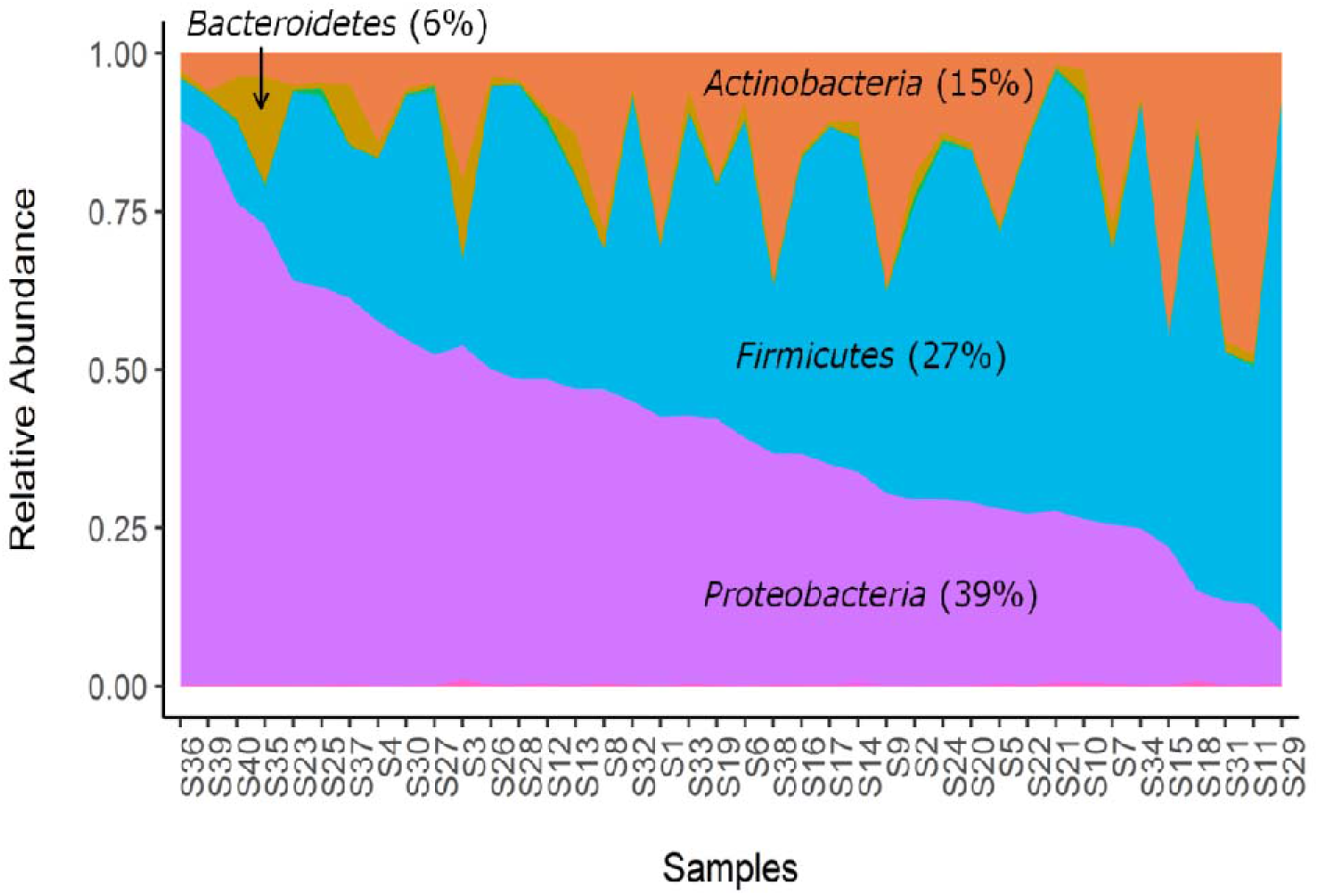
Stackplot of the phyla distribution across the 40 sperm samples.

**Table 1.**
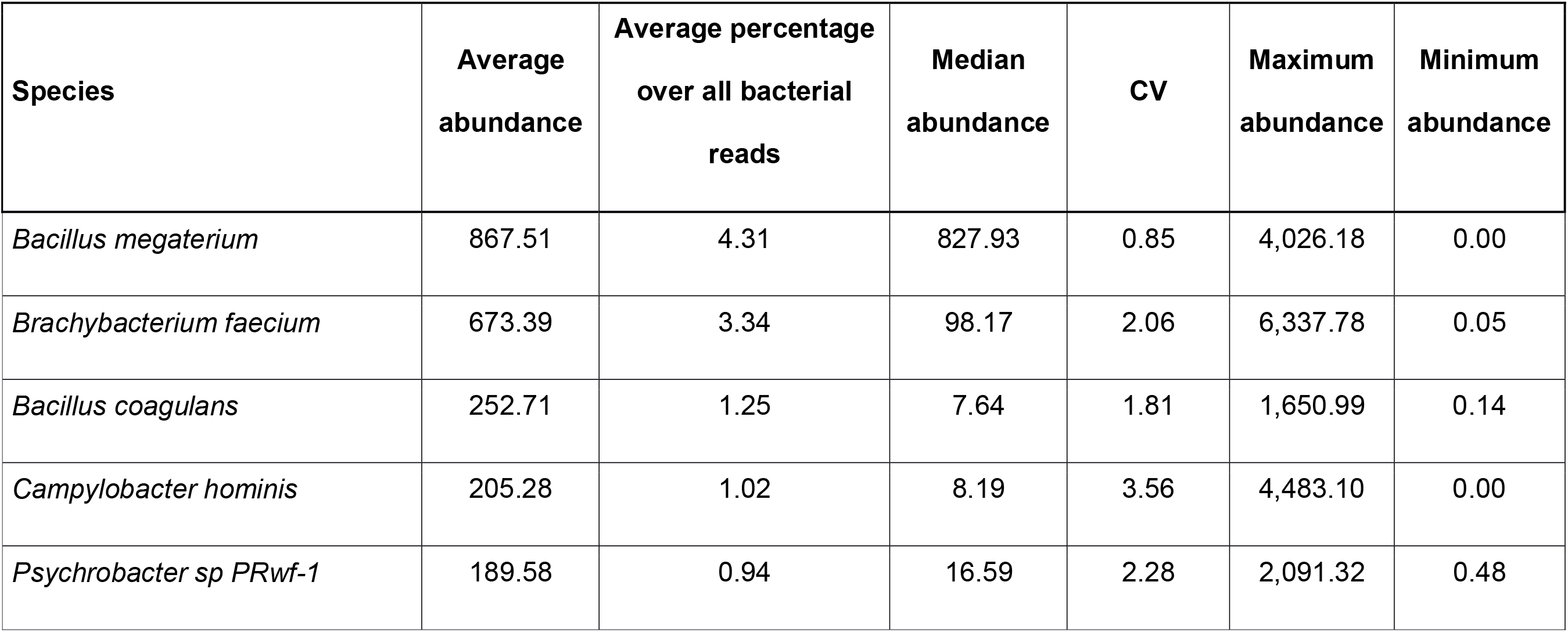
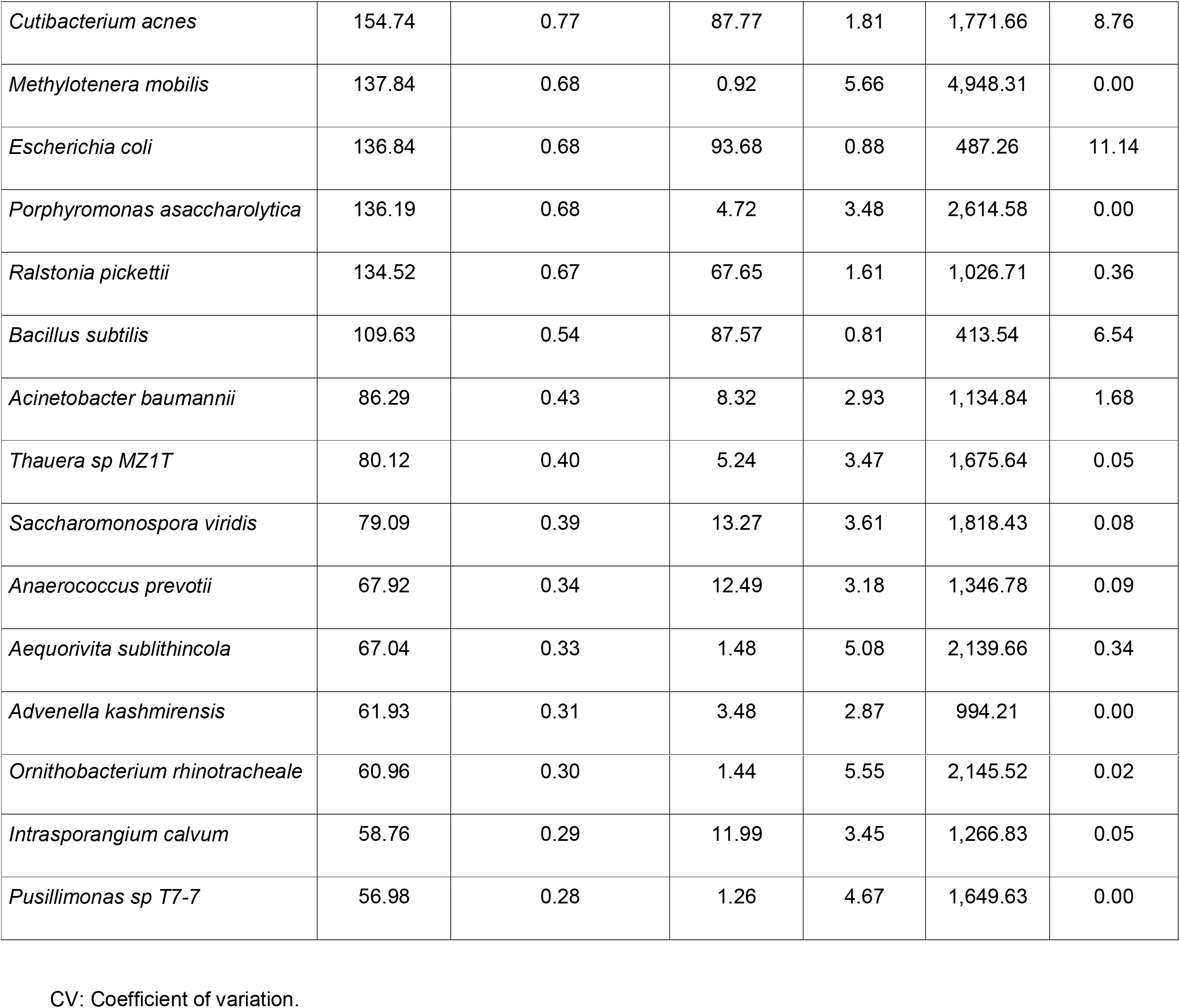
List of the 20 most abundant bacteria in the boar sperm.

### Boar sperm safety: pathogens and antibiotic resistance genes

We found 11 potentially pathogenic species of bacteria with average abundance ≥ 1 CPM and representing more than 0.001% of the total bacterial read counts, according to the Professional Pig Community pig333 site (www.pig333.com/pig-diseases) but only 7 displayed CPM > 5. These were, in this order: *Escherichia coli, Clostridioides difficile, Clostridium perfringens, Clostridium botulinum, Mycobacterium tuberculosis, Mycoplasma hyopneumoniae* and *Campylobacter jejuni* (Table 2). With the exception of *E. coli* and *C. difficile*, which ranked 8^th^ and 22^nd^ in the list of most abundant bacteria species, with 137 and 50 CPM, respectively, these potential bugs were in general displaying low abundance in our samples (Table 2). While nearly all the samples contained at least traces of these bacteria, *M. tuberculosis* was only present in 6 samples and it presented moderate abundances (between 28 and 84 CPMs) in all of them (Table S2).

**Table 2.** List of potential pathogens and antimicrobial resistance genes identified in pig sperm.

| Potential pathogen species | Average abundance | Median abundance | CV | Maximum abundance | Minimum abundance | Disease / health condition |
| --- | --- | --- | --- | --- | --- | --- |
| *Escherichia coli* | 136.84 | 93.68 | 0.88 | 487.26 | 11.14 | Diarrhoea and high mortality in piglets |
| *Clostridioides difficile* | 49.77 | 16.74 | 1.76 | 338.49 | 0.88 | Diarrhoea in piglets |
| *Clostridium perfringens* | 14.05 | 5.60 | 1.47 | 95.99 | 0.47 | Chronic or acute enteritis in piglets. Sometimes also gangrene and sudden death in adults |
| *Clostridium botulinum* | 7.67 | 2.92 | 1.71 | 69.73 | 0.33 | Toxins produced by this bacteria cause a progressive flaccid paralysis, but pigs are very resistant to the toxin |
| *Mycobacterium tuberculosis* | 7.46 | 0.00 | 2.60 | 84.45 | 0.00 | Tuberculosis |
| *Mycoplasma hyopneumoniae* | 6.24 | 5.48 | 0.78 | 23.39 | 0.82 | Enzootic pneumonia |
| *Campylobacter jejuni* | 5.86 | 0.05 | 3.11 | 90.93 | 0.00 | Clinical signs are not always present but can cause a watery diarrhea with mucous and blood. Also food-borne illness in humans |
| *Erysipelothrix rhusiopathiae* | 3.91 | 0.64 | 2.43 | 42.22 | 0.00 | Erysipela: skin lesion and arthritis |
| *[Haemophilus] parasuis* | 3.00 | 0.12 | 3.81 | 67.62 | 0.00 | Glässer disease: polyserosistis and sporadic arthritis |
| *Streptococcus suis* | 2.42 | 0.60 | 2.35 | 29.64 | 0.05 | Streptococcal infection with pneumonia, septicemia, arthritis, etc. Zoonotic potential |
| *Listeria monocytogenes* | 2.21 | 0.97 | 1.70 | 21.40 | 0.02 | Rare systemic bacterial septicemia |
| **Potential antibiotic resistant gene** |  |  |  |  |  |  |
| ARO:3003497_Neisseria_meningitidis_16S_rRNA_mutation_spectinomycin | 27.85 | 21.95 | 1.11 | 189.86 | 2.31 |  |
| ARO:3004058_Staphylococcus_aureus_23S_rRNA_with_mutation_linezolid | 125.07 | 103.89 | 0.78 | 400.27 | 5.16 |  |
| ARO:3004150_Escherichia_coli_23S_rRNA_with_mutation_chloramphenicol | 316.72 | 198.07 | 1.01 | 1634.58 | 113.99 |  |

CV: Coefficient of variation.

We also searched for ARG with average CPM ≥ 1 and found 3 candidates, including ARO:3003497, *Neisseria meningitidis* 16S rRNA mutation conferring resistance to spectinomycin; ARO:3004058, *Staphylococcus aureus* 23S rRNA with mutation conferring resistance to linezolid and ARO:3004150, *E. coli* 23S rRNA with mutation conferring resistance to chloramphenicol. Moreover, all the samples presented CPM ≥ 1 for these 3 ARGs (Table 2).

### Relationship between the sperm microbiome and semen quality

To identify potential relationships between bacterial abundances and semen quality we employed Link-HD (Zingaretti et al., 2019), a recently developed tool based on STATIS methodology to integrate heterogeneous datasets. This approach analyzes different types of variables measured on the same samples, here bacterial abundance and semen quality phenotypes. To the end, the tool turns each raw data into cross-product matrix, computed on the distances between samples, which are then combined in a common configuration named compromise. A classical Principal Component Analysis (PCA) decomposes the compromise variance into orthogonal components and data structure can be easily recovered using standard clustering techniques. In this study, the samples were clustered into categories according to their microbiome and their semen quality. After quality controls, we included the 733 bacterial features (from phyla to species in Table S2) and 7 semen quality traits (Table S3). Link-HD structured the purified ejaculates into 2 clusters with 30 (cluster 1) and 10 (cluster 2) samples each (Fig. 3 and Table S4). The analysis also recovers the contribution of each feature into the common structure, which facilitates the interpretability of the results. We found that the 7 semen traits and 67 of the 733 bacterial features associated with the whole-compromise structure (Table S5).

**Figure 3.**
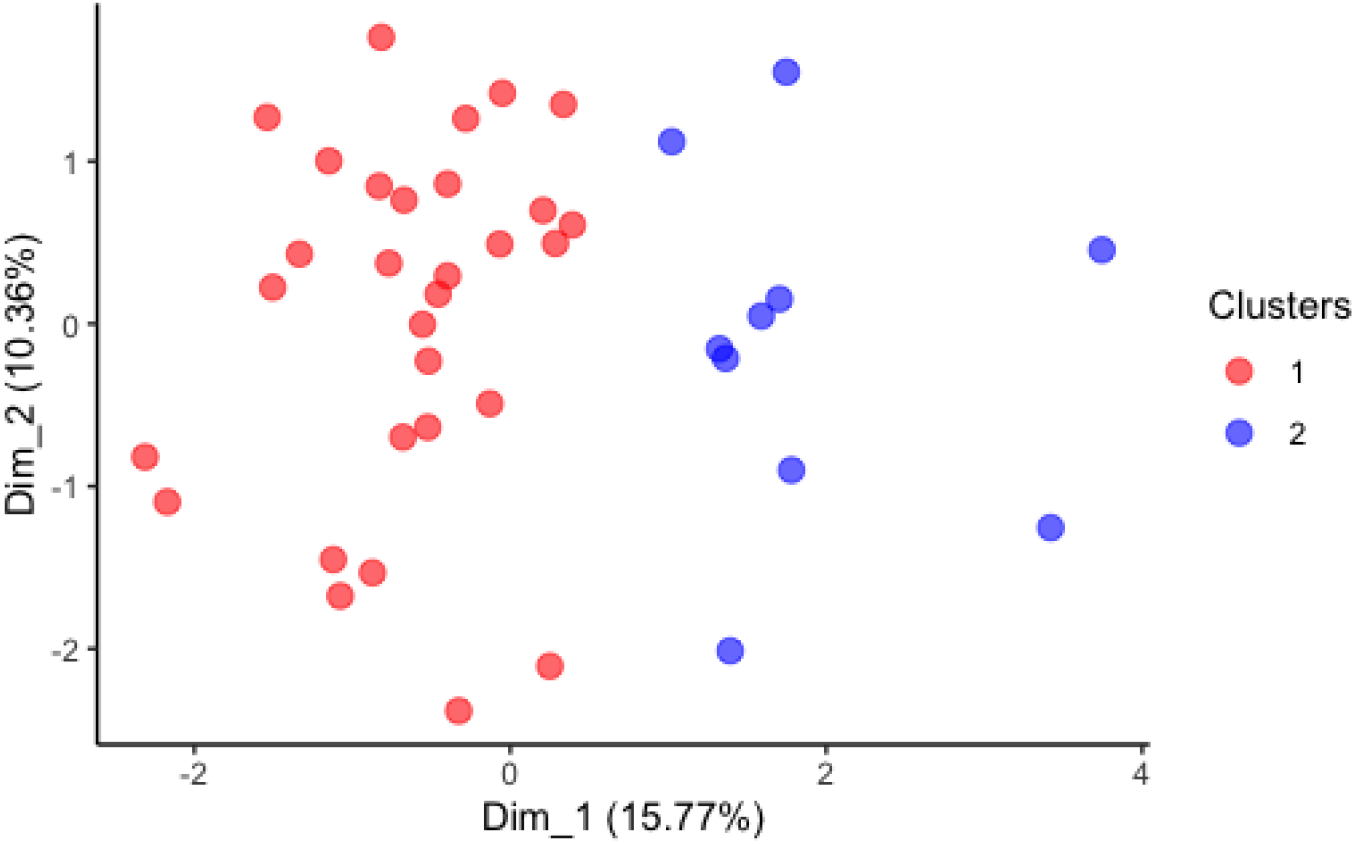
Data structure from compromise configuration after applying a clustering using standard k-means with Link-HD.

We then compared the distribution of these 7 phenotypes and 67 bacterial features in each cluster. The 2 categories showed statistically significant differences for 5 traits. MT_5 and MT_90 did not differ between both groups (Table 3). The feature abundances between the 2 clusters were compared. Thirteen bacterial features resulted in nominal significant differences between clusters (Table 4). These included the genus *Acinetobacter*, *Stenotrophomonas* and *Rhodobacter* (Table 4).

**Table 3.** List of phenotypes displaying significant differences between the 2 clusters distinguishing both groups.

| Parameter | VIAB_5 | VIAB_90 | ACRO_5 | ACRO_90 | ORT | MT_5 | MT_90 |
| --- | --- | --- | --- | --- | --- | --- | --- |
| Average cluster 1 | 0.39 | 3.78 | -0.71 | -2.00 | 3.81 | -1.28 | -0.58 |
| Average cluster 2 | -3.92 | -14.59 | 5.16 | 9.31 | -10.24 | 0.94 | -3.06 |
| Average all | -0.69 | -0.81 | 0.76 | 0.83 | 0.30 | -0.72 | -1.20 |
| SD cluster 1 | 3.50 | 6.38 | 3.37 | 3.62 | 4.55 | 17.48 | 21.12 |
| SD cluster 2 | 3.78 | 11.02 | 5.28 | 9.53 | 8.61 | 13.81 | 26.37 |
| Total SD | 4.00 | 11.10 | 4.63 | 7.44 | 8.40 | 16.50 | 22.21 |
| Mann Whitney U test (P-value) | 0.003 | 1.1E-05 | 0.004 | 1.5E-04 | 2.3E-05 | 0.863 | 1.000 |

SD: Standard deviation; VIAB\_5: percentage of viable sperm cells after 5 minutes of incubation at 37 °C; VIAB\_90: percentage of viable sperm cells after 90 min incubation at 37 °C; ACRO\_5: percentage of cells with abnormal acrosomes after the 5 min; ACRO\_90: 90 min incubation; ORT, percentage of viable cells after an osmotic stress (Osmotic Resistance Test); MT\_5: percentage of motile cells after 5 min; MT\_90: 90 min incubation.

**Table 4.**
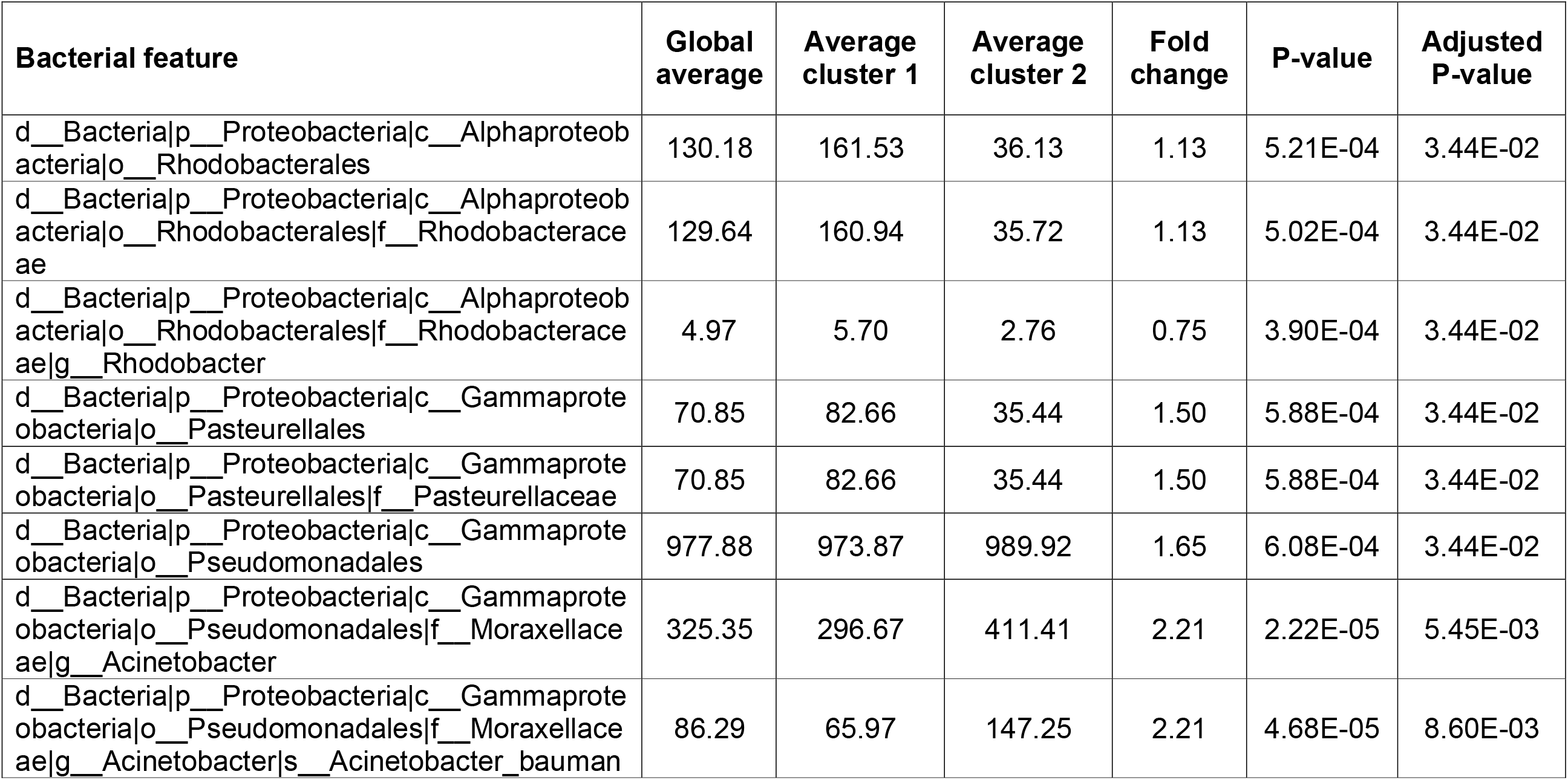
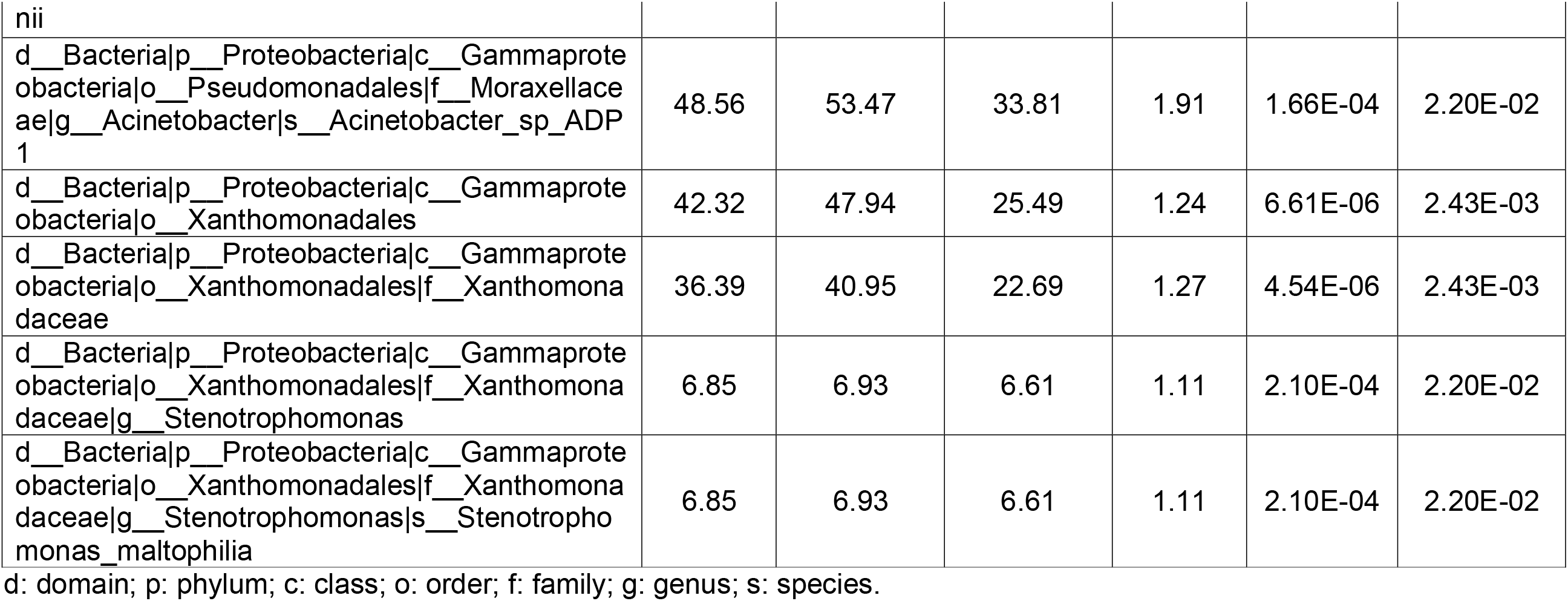
List of bacteria displaying significant differences between clusters.

## Discussion

### Technical considerations

Bacterial composition and abundance in different animal compartments has been proven to affect a plethora of phenotypes in humans and domestic animals. This has become more obvious with the recent development of high throughput sequencing tools that allow the systematic characterization of microbiomes (Pennisi, 2007). In 2014, Weng et al. (Weng et al., 2014) identified a particular microbiome composition in human ejaculated sperm and related the abundance of several bacteria, most remarkably *Lactobacillus* and *Prevotella*, to semen quality. We carried RNA-seq on the extended sperm from 40 pigs after purification of the samples to remove somatic cells and bacteria with Bovipure^™^ with the aim to characterize the boar semen transcriptome in relation to sperm quality. After mapping the RNA-seq reads to the pig genome (Sscrofa11.1), between 9 and 31% of the reads remained unmapped (Table S1). We hypothesized that a proportion of the unmapped sequences could correspond to bacterial transcripts and aligned these reads to microbiome genomes. We identified a rich population of bacteria with a diverse abundance profile between the ejaculates. Despite the fact that we processed extended sperm containing antibiotics and that we treated these samples to remove micro-organisms, we found evidences of bacterial presence in their sequenced RNAs. This indicates that the extender does not eliminate all the bacteria present in the ejaculate. It is even possible that these antibiotics promote a positive selection for the antibiotic resistant bacteria. In fact, we observed the presence of 3 ARGs that confer resistance to spectinomycin, linezolid and chloramphenicol. The results also suggest that a large amount of bacteria may be present in the sperm doses used in artificial insemination.

The bioinformatics tool Kraken (Wood and Salzberg, 2014) was designed to study meta-genomes and quantify the abundance of bacteria based in their DNA. Kraken has been also used to characterize the sperm microbiome using RNA-seq datasets in human (Swanson et al., 2020). While meta-genomics strictly focuses on the abundance of bacterial specimens, meta-transcriptomics informs on the expression of their genes and thus the function and activity of these micro-organisms in the sample. Our data provides a quantification of each bacterium based in the overall expression of their transcripts which accounts for both the bacterial abundance and their gene expression activity and have the additional advantage to account for active microorganisms. In other words, we cannot state without uncertainty whether one bacterium is more abundant than another in one sample but we can assume that this is the most likely scenario as in part, our measures are reflecting these abundances. For this reason and to ease the message provided in this manuscript, we have referred to bacterial abundance throughout the article.

### Sperm microbial composition

According to our data, the boar sperm microbiome differed from the profiles obtained on porcine gut where the most abundant phyla include *Bacteroidetes* and *Firmicutes* and the predominant genus are *Prevotella* and *Roseburia* (Xiao et al., 2016). On the contrary, our data highlights that in the porcine and in human sperm, the 4 most abundant phyla are coincident (Swanson et al., 2020). Moreover, 11 of the 20 most abundant genera in boar and human sperm were shared in both species. In human sperm, the most abundant bacteria were members of *Actinobacteria* (*Corynebacterium*), *Bacteroidetes* (*Prevotella*), *Firmicutes* (*Lactobacillus*, *Streptococcus*, *Staphylococcus*, *Planococcaceae*, *Finegoldia*), and *Proteobacteria* (*Haemophilus*, *Burkholderia*) (Baud et al., 2019). The differences between the porcine and the human ejaculates could be attributed to multiple technical (e.g., the selection of antibiotics in extender and the removal of bacteria during the purification of the samples), environmental and biological causes. Although boar studs are kept in high hygienic conditions, pigs are in closer contact with surfaces, soil, faeces and water and are thus more exposed to environmental contaminants than humans.

The most abundant bacteria in the boar sperm are mostly environmental (*B. megaterium* (Vary, 1994), *B. faecium* (Collins et al., 1988), *R. pickettii* (Labarca et al., 1999)) and faecal (*C. hominis* (Lawson et al., 1998) and *E. coli*). This suggests that these bacteria have contaminated sperm after ejaculation. *C. acnes* typically colonizes the human skin (Brüggemann et al., 2004) but can be also found in other compartments including the gastrointestinal tract (Perry and Lambert, 2011). Interestingly, *B. subtilis*, a probiotic added in the pig feedstuff and allowed in the European Union Register of Feed Additives, appeared as the 11^th^ most abundant bacteria in the boar sperm (Table 1), again suggesting that it contaminated sperm after ejaculation.

Arkfen and co-authors (Arfken et al., 2015) analysed the airborne microbiome of hog farms and found a similar composition of bacterial phyla as the one described in our study. Moreover, our data is in line with the latest comprehensive study carried by Schulze et al. (Schulze et al., 2015) which indicated that the major sources of contamination were of non-animal origin and were mostly attributed to sinks and drains but also to other utensils and the skin flora of working staff. This further supports the hypothesis that most of the bacteria present in the pig sperm are from environmental contamination.

Three ejaculates showed a much higher bacterial abundance when compared to the average in all the samples (Fig. 1). Although we don’t know the causes, these elevated values of bacterial reads might have been caused by accidental contamination of the ejaculate with particularly large chunks of environmental debris present for example in the boar’s prepuce or other surfaces.

### Pathogens and anti-microbial resistances

We found several potential pathogens (Table 2) as included in the Professional Pig Community pig333 site (www.pig333.com/pig-diseases). Some serotypes of these bacteria have been linked to diarrhea (*E. coli*, *C. difficile*, *C. jejuni*), acute enteritis (*C. perfringens*), botulism (*C. botulinum*), tuberculosis (*M. tuberculosis*) and enzootic pneumonia (*M. hyopneumoniae*) in swine. While 4 of the 5 most abundant potential pathogens showed a continuous pattern of abundance across samples, *M. tuberculosis* was only present in 6 samples, all with moderate abundances (CPM between 28 and 84). This quasi bi-modal distribution cannot be explained by factors controlled in our study as these 6 pigs came from different farms, were of varying ages, were collected at different seasons of the year and there was thus no apparent link between these animals. The pathogenic potential of these bacterial species varies across strains depending on the presence of virulence factors and toxin production. Notwithstanding, our analysis does not allow concluding that any of the specimens identified in this study are pathogenic as the analysis did not have the power and specificity to detect the genes to discriminate between these serotypes.

In animal production systems, extended sperm is distributed to multiple farms and geographical locations and despite the fact that it is mixed with antibiotics, some bacteria remain in these ejaculates. Moreover, before they are inseminated into the sow, extended sperm doses will remain at 17 °C in average up to few days, thus potentially allowing the selective growth of bacteria carrying ARGs. Therefore, ejaculates might be an important source and vehicle to disseminate these bacteria to other farms and animals.

The evaluation of the vaginal microbiome in sows inseminated with these doses should be evaluated to determine how the sperm microbiome modulates the vaginal counterpart and how it impacts on the sow’s health and fertility and the extent to which ARGs and pathogens are transmitted through artificial insemination.

We identified 3 ARGs that were present at CPM ≥ 1 in all the ejaculates (Table 2). These ARGs were point mutations in bacterial ribosomal RNA genes. The most abundant ARG potentially conferred resistance to *E. coli* to chloramphenicol, a broad spectrum antibiotic predominantly active against gram negative bacteria used in human medicine but not authorized by the European Union for use in livestock. However, this antimicrobial can be synthesized by soil bacteria and it may thus be present in farms thereby allowing the generation of ARGs against it. Our results suggest a scarce presence of ARGs in our porcine sperm samples. The ejaculates were diluted with a commercial semen extender that contains the antibiotics apramycin, cephalosporin and gentamicin but no ARGs were found against these 3 antibiotics.

The 3 bacteria involved in these presumable ARGs (*E. coli*, *N. meningitidis* and *S. aureus*) were detected in our study but linear regression showed that their abundances did not relate with the expression levels of their cognate ARGs (Fig. S1). The only exception is for *N. meningitidis* and the ARG for Specinomycin (R^2^ = 0.74), but this is largely due to one influential outlier ejaculate for which the abundance of both, these bacteria and ARG were remarkably elevated (Fig. S1). This indicates that not all the bacteria of these species carry the same load of ARG in each sample.

These results have to be taken as indicative as in this study we cannot conclude whether these abundances in CPM are large or modest. Moreover, the antimicrobial activity of these ARGs cannot be granted and should be confirmed with a microbiological analysis by growing the bacteria in media containing the target antibiotic using, for example, the agar disc diffusion method. The appearance and spread of antibiotic resistant bacteria is one of the biggest present and future challenges of our society due to their impact on animal, environmental and human health. These resistances are growing and jeopardizing the treatments to fight bacterial infections. Recently, a resistance against collistin, a latest resource antibiotic used to eliminate *C. difficile* from hospitals appeared in a pig farm in China (Collins and Riley, 2018). This ARG disseminated quickly and reached human hospitals and livestock farms throughout the world. *C. difficile* is a bacteria difficult to treat that threatens some patients and have fatal consequences (Collins and Riley, 2018). In the framework of One Health, actors from multiple sectors (animal breeders, researchers and policy makers) are seriously studying to reduce the use of antibiotics to decrease the rate of development of ARGs. In this sense, as happens with the dissemination of pathogen bacteria, ejaculates can be responsible for a significant proportion of the dissemination of ARGs between farms.

### Relationship between the sperm microbiome and semen quality

Motivated by the results from Weng et al. that found a relationship between male fertility and the microbiome of the human sperm (Weng et al., 2014) we sought to assess whether there was a relation between the sperm microbiome and semen quality. Studies in pig have found detrimental associations between *E. coli* and sperm cell agglutination and reduced fertility (Maroto Martin et al., 2010), *C. perfringens* and diminished sperm motility and viability (Sepúlveda et al., 2013) and *Pseudomonas aureginosa* and decreased longevity and fertilizing ability (Sepúlveda et al., 2014).

The microbiome is a complex system of microbial communities and its genomic characterization generates compositional and sparse data. Thus, we decided to treat the sperm bacterial communities holistically when assessing its relation to semen quality instead of analyzing the individual impact of each bacterial entity. For this, we used an integrative approach that considers simultaneously the ejaculate bacterial composition and semen quality. This analysis led to the identification of two clusters with 30 and 10 samples each. Five traits and 13 bacterial features showed significant differences between the two clusters (Table 3 and Table 4). Semen quality was worst for cluster 2 (Table 3). Larger values for VIAB and ORT indicate more robust and viable semen whilst higher values for ACRO indicate a larger proportion of spermatozoa with deficient acrosomes.

The fact that this analysis identified two clear categories based simultaneously on their semen quality and microbiome indicates that the two are related. Interestingly, VIAB_90 and ACRO_90 displayed stronger differences between the two groups than VIAB_5 and ACRO_5. This suggests that the long incubation favored the proliferation of bacteria and this lead to a stronger bacterial impact on the phenotype. In farm conditions, extended ejaculates are kept at 17 °C until they are used for insemination. Most doses are used within 48 h after ejaculation but some may be kept up to 6 days. The impact of these conditions in the microbiome could be strong and it should be explored. The 13 bacterial features showing differences between the two clusters included the genera *Acinetobacter*, *Stenotrophomonas* and *Rhodobacter* (Table 4). *Acinetobacter* are typically found in soil, although *A. baumannii* can be also detected in hospitals where it can act as an opportunistic pathogen affecting immunocompromised patients. One study on human semen from Kiessling et al. (Kiessling et al., 2008) identified *Acinetobacter* bacteria in some of the semen samples that they evaluated (Kiessling et al., 2008). An *in vitro* study on rabbit sperm cultured under the presence of *A. baumannii* showed that the motility of the spermatozoa was negatively affected by the presence of this bacterium (Tvrdá et al., 2018). A study on boar sperm found *A. iwoffi* in some samples and that the presence of this bacteria was associated to higher production of Reactive Oxidative Species and lipid peroxidation thus potentially altering some semen quality features not measured in that study (Ďuračka and Tvrda, 2018). *Stenotrophomonas* are also typically found in soil and plants and some (including *S. maltophila*) can be opportunistic pathogens in humans. In swine, it has been previously detected in sperm (Althouse and Lu, 2005). A case report on a dog with conception failure and positive for *S. malthophila*, linked this bacteria with semen quality (Domoslawska et al., 2017). Finally, the genus *Rhodobacter* includes several species with a diverse range of energy-based metabolism but has not been previously found in sperm nor linked to sperm quality. This genus can be found in varied habitats including pig manure (Yang et al., 2011).

## Conclusions

In conclusion, we have identified a large and varied population of bacteria contaminating the boar’s extended sperm, including a small proportion of potential pathogens and ARGs. Moreover, some of these bacteria, such as *Acinetobacter*, *Stenotrophomonas* and *Rhodobacter*, seem to be related to semen quality. This is of high relevance for two main reasons. First, these bacteria may affect sperm quality and male fertility. Second, since ejaculates are widely distributed across farms, they might be major disseminators of these microbes and ARGs. Thus, the microbial composition in the sperm of swine and other livestock species needs to be studied more profoundly. Moreover, we anticipate that in a not too distant future, the systematic microbiome analysis of semen ejaculates to identify the samples that contain potential pathogens will become common practice. At present, high throughput sequencing is still an expensive technology and this makes its routine application to assess semen quality in swine unfeasible. However, these costs are expected to keep decreasing in the years to come. This drop in sequencing costs should allow the systematic implementation of metagenomics to routinely assess the presence of pathogens and ARGs in the boar sperm.

## Acknowledgements

We thank Sam Balasch (grup Gepork S.A.) and Craig Lewis (Genus - PIC) for providing the blood and sperm samples. This work was supported by the Spanish Ministry of Economy and Competitiveness (MINECO) under grant AGL2013-44978-R and grant AGL2017-86946-R and by the CERCA Programme/Generalitat de Catalunya. AGL2017-86946-R was also funded by the Spanish State Research Agency (AEI) and the European Regional Development Fund (ERDF). We thank the Agency for Management of University and Research Grants (AGAUR) of the Generalitat de Catalunya (Grant Numbers 2014 SGR 1528 and 2017 SGR 01060). We also acknowledge the support of the Spanish Ministry of Economy and Competitivity for the Center of Excellence Severo Ochoa 2016–2019 (Grant Number SEV-2015-0533) grant awarded to the Centre for Research in Agricultural Genomics (CRAG). MG acknowledges a Ph.D. studentship from MINECO (Grant Number BES-2014-070560). YRC was funded by Marie Skłodowska-Curie grant (P-Sphere) agreement No 6655919 (EU). LMZ is recipient of a Ph.D. grant associated with the SEV-2015-0533 award to CRAG.

## Availability of data

The datasets generated and analysed are available at NCBI’s BioProject PRJNA520978.

## Supporting information

**Supplementary Table S1.** RNA-seq statistics for each of the 40 samples.

SD: Standard Deviation.

**Supplementary Table S2.** Full list of bacteria taxa and its abundance in the 40 samples.

CPM: Counts per Million reads.

**Supplementary Table S3.** Phenotypic values for the 7 semen quality traits for each of the 40 samples.

**Supplementary Table S4.** Detail of the samples belonging to each Link_HD cluster.

**Supplementary Table S5.** Detail of the traits and bacterial features contributing to the Link-HD compromise.

**Supplementary Figure S1.** Linear regression plots of the abundance of the ARGs and their related bacterial species.

## References

Althouse, G. C. & Lu, K. G. (2005). Bacteriospermia in extended porcine semen. Theriogenology 63, 573–84.

Arfken, A. M., Song, B. & Sung, J.-S. (2015). Comparison of airborne bacterial communities from a hog farm and spray field. Journal of Microbiology and Biotechnology 25, 709–17.

Bao, W., Kojima, K. K. & Kohany, O. (2015). Repbase Update, a database of eukaryotic repetitive elements. Mobile DNA 6, 11.

Baud, D., Pattaroni, C., Vulliemoz, N., Castella, V., Marsland, B. J. & Stojanov, M. (2019). Sperm Microbiota and Its Impact on Semen Parameters. Frontiers in Microbiology 10, 1–9.

Bolger, A. M., Lohse, M. & Usadel, B. (2014). Trimmomatic: A flexible trimmer for Illumina sequence data. Bioinformatics 30, 2114–20.

Brüggemann, H., Henne, A., Hoster, F., Liesegang, H., Wiezer, A., Strittmatter, A., Hujer, S. & Du, P. (2004). The Complete genome of Propionibacterium Acnes, a Commensal of Human Skin. Science 305, 671–4.

Camacho, C., Coulouris, G., Avagyan, V., Ma, N., Papadopoulos, J., Bealer, K. & Madden, T. L. (2009). BLAST+: Architecture and applications. BMC Bioinformatics 10, 1–9.

Collins, D. A. & Riley, T. V. (2018). Clostridium difficile in Asia: Opportunities for One Health Management. Tropical Medicine and Infectious Disease 4, 7.

Collins, M. D., Brown, J. & Jones, D. (1988). Brachybacterium faecium gen. nov., sp. nov., a Coryneform Bacterium from Poultry Deep Litter. International Journal of Systematic Bacteriology 38, 45–8.

Domoslawska, A., Zdunczyk, S., Jurczak, A. & Janowski, T. (2017). Stenotrophomonas maltophilia isolated from prostatic fluid as an infertility factor in a male dog. Andrologia 49, e12769.

Ďuračka, M. & Tvrda, E. (2018). The presence of bacterial species in boar semen and their impact on the sperm quality and oxidative balance. Journal of Animal Science 96, 501.

Gòdia, M., Estill, M., Castelló, A., Balasch, S., Rodríguez-Gil, J. E., Krawetz, S. A., Sánchez, A. & Clop, A. (2019). A RNA-Seq Analysis to Describe the Boar Sperm Transcriptome and Its Seasonal Changes. Front Genet 10, 299.

Gòdia, M., Mayer, F. Q., Nafissi, J., Castelló, A., Rodríguez-Gil, J. E., Sánchez, A. & Clop, A. (2018). A technical assessment of the porcine ejaculated spermatozoa for a sperm-specific RNA-seq analysis. Systems Biology in Reproductive Medicine 64, 291–303.

Hou, D., Zhou, X., Zhong, X., Settles, M., Herring, J., Wang, L., Abdo, Z., Forney, L. J. & Xu, C. (2013). Microbiota of the seminal fluid from healthy and infertile men. Fertility and Sterility 100, 1261–9.

Jia, B., Raphenya, A. R., Alcock, B., Waglechner, N., Guo, P., Tsang, K. K., Lago, B. A., Dave, B. M., Pereira, S., Sharma, A. N., Doshi, S., Courtot, M., Lo, R., Williams, L. E., Frye, J. G., Elsayegh, T., Sardar, D., Westman, E. L., Pawlowski, A. C., Johnson, T. A., Brinkman, F. S. L., Wright, G. D. & Mcarthur, A. G. (2017). CARD 2017: Expansion and model-centric curation of the comprehensive antibiotic resistance database. Nucleic Acids Research 45, D566–D73.

Kiessling, A. A., Desmarais, B. M., Yin, H. Z., Loverde, J. & Eyre, R. C. (2008). Detection and identification of bacterial DNA in semen. Fertility and Sterility 90, 1744–56.

Kim, D., Langmead, B. & Salzberg, S. L. (2015). HISAT: a fast spliced aligner with low memory requirements. Nature Methods 12, 357–60.

Labarca, J. A., Trick, W. E., Peterson, C. L., Carson, L. A., Holt, S. C., Arduino, M. J., Meylan, M., Mascola, L. & Jarvis, W. R. (1999). A multistate nosocomial outbreak of Ralstonia pickettii colonization associated with an intrinsically contaminated respiratory care solution. Clinical infectious diseases: an official publication of the Infectious Diseases Society of America 29, 1281–6.

Lawson, A. J., Linton, D. & Stanley, J. (1998). 16S rRNA gene sequences of ‘Candidatus Campylobacter hominis’, a novel uncultivated species, are found in the gastrointestinal tract of healthy humans. Microbiology 144, 2063–71.

Mahboubi, M. A., Carmody, L. A., Foster, B. K., Kalikin, L. M., Vandevanter, D. R. & Lipuma, J. (2016). Cystic Fibrosis Respiratory Specimens. Journal of clinical microbiology 54, 613–9.

Mändar, R., Punab, M., Borovkova, N., Lapp, E., Kiiker, R., Korrovits, P., Metspalu, A., Krjutškov, K., Nlvak, H., Preem, J. K., Oopkaup, K., Salumets, A. & Truu, J. (2015). Complementary seminovaginal microbiome in couples. Research in Microbiology 166, 440–7.

Maroto Martin, L. O., Munoz, E. C., De Cupere, F., Van Driessche, E., Echemendia-Blanco, D., Rodriguez, J. M. & Beeckmans, S. (2010). Bacterial contamination of boar semen affects the litter size. Anim Reprod Sci 120, 95–104.

Moreno, I., Codoñer, F. M., Vilella, F., Valbuena, D., Martinez-Blanch, J. F., Jimenez-Almazán, J., Alonso, R., Alamá, P., Remohí, J., Pellicer, A., Ramon, D. & Simon, C. (2016). Evidence that the endometrial microbiota has an effect on implantation success or failure. American Journal of Obstetrics and Gynecology 215, 684–703.

Paulson, J. N., Pop, M. & Bravo, H. C. (2013). metagenomeSeq: Statistical analysis for sparse high-throughput sequencing. Bioconductor package.

Pennisi, E. (2007). METAGENOMICS: Massive Microbial Sequence Project Proposed. Science 315, 1781a.

Perry, A. & Lambert, P. (2011). Propionibacterium acnes: infection beyond the skin. Expert Review of Anti-infective Therapy 9, 1149–56.

R Core Team. 2017. R: A language and environment for statistical computing.R Foundation for Statistical Computing, Vienna, Austria.

Schroeder, B. O. & Bäckhed, F. (2016). Signals from the gut microbiota to distant organs in physiology and disease. Nature Medicine 22, 1079–89.

Schulze, M., Ammon, C., Rüdiger, K., Jung, M. & Grobbel, M. (2015). Analysis of hygienic critical control points in boar semen production. Theriogenology 83, 430–7.

Schulze, M., Schäfer, J., Simmet, C., Jung, M. & Gabler, C. (2018). Detection and characterization of Lactobacillus spp. In the porcine seminal plasma and their influence on boar semen quality. PLoS ONE 13, 1–16.

Sepúlveda, L., Bussalleu, E., Yeste, M. & Bonet, S. (2014). Effects of different concentrations of Pseudomonas aeruginosa on boar sperm quality. Animal Reproduction Science 150, 96–106.

Sepúlveda, L., Bussalleu, E., Yeste, M. & Bonet, S. (2016). Effect of Pseudomonas aeruginosa on sperm capacitation and protein phosphorylation of boar spermatozoa. Theriogenology 85, 1421–31.

Sepúlveda, L., Bussalleu, E., Yeste, M., Torner, E. & Bonet, S. (2013). How do different concentrations of Clostridium perfringens affect the quality of extended boar spermatozoa? Animal Reproduction Science 140, 83–91.

Swanson, G. M., Moskovtsev, S., Librach, C., Pilsner, J. R., Goodrich, R. & Krawetz, S. A. (2020). What human sperm RNA-Seq tells us about the microbiome. Journal of Assisted Reproduction and Genetics 37, 359–68.

Tvrdá, E., Ďuračka, M., Kántor, A., Halenár, M. & Hleba, L. (2018). In *vitro* effects of *Acinetobacter Baumannii* and selected natural biomolecules on rabbit spermatozoa motility. AGROFOR International Journal 3, 54–63.

Úbeda, J. L., Ausejo, R., Dahmani, Y., Falceto, M. V., Usan, A., Malo, C. & Perez-Martinez, F. C. (2013). Adverse effects of members of the Enterobacteriaceae family on boar sperm quality. Theriogenology 80, 565–70.

Vary, P. S. (1994). Prime Time for Bacillus megaterium. Microbiology 140, 1001–13.

Weng, S. L., Chiu, C. M., Lin, F. M., Huang, W. C., Liang, C., Yang, T., Yang, T. L., Liu, C. Y., Wu, W. Y., Chang, Y. A., Chang, T. H. & Huang, H. D. (2014). Bacterial communities in semen from men of infertile couples: Metagenomic sequencing reveals relationships of seminal microbiota to semen quality. PLoS ONE 9, e110152.

Wood, D. E. & Salzberg, S. L. (2014). Kraken: ultrafast metagenomic sequence classification using exact alignments. Genome Biology 15, R46.

Xiao, L., Estellé, J., Kiilerich, P., Ramayo-Caldas, Y., Xia, Z., Feng, Q., Liang, S., Pedersen, A. O., Kjeldsen, N. J., Liu, C., Maguin, E., Dore, J., Pons, N., Le Chatelier, E., Prifti, E., Li, J., Jia, H., Liu, X., Xu, X., Ehrlich, S. D., Madsen, L., Kristiansen, K., Rogel-Gaillard, C. & Wang, J. (2016). A reference gene catalogue of the pig gut microbiome. Nat Microbiol 1, 16161.

Yang, Y. Y., Pereyra, L. P., Young, R. B., Reardon, K. F. & Borch, T. (2011). Testosterone-Mineralizing Culture Enriched from Swine Manure: Characterization of Degradation Pathways and Microbial Community Composition. Environmental Science& Technology 45, 6879–86.

Zingaretti, L. M., Renand, G., Morgavi, D. P. & Ramayo-Caldas, Y. (2019). Link-HD: a versatile framework to explore and integrate heterogeneous microbial communities. Bioinformatics, 1–2.

