## Supplementary figures and images for "A RNA-seq characterization of the porcine sperm microbiome"

### Supplementary Figure S1

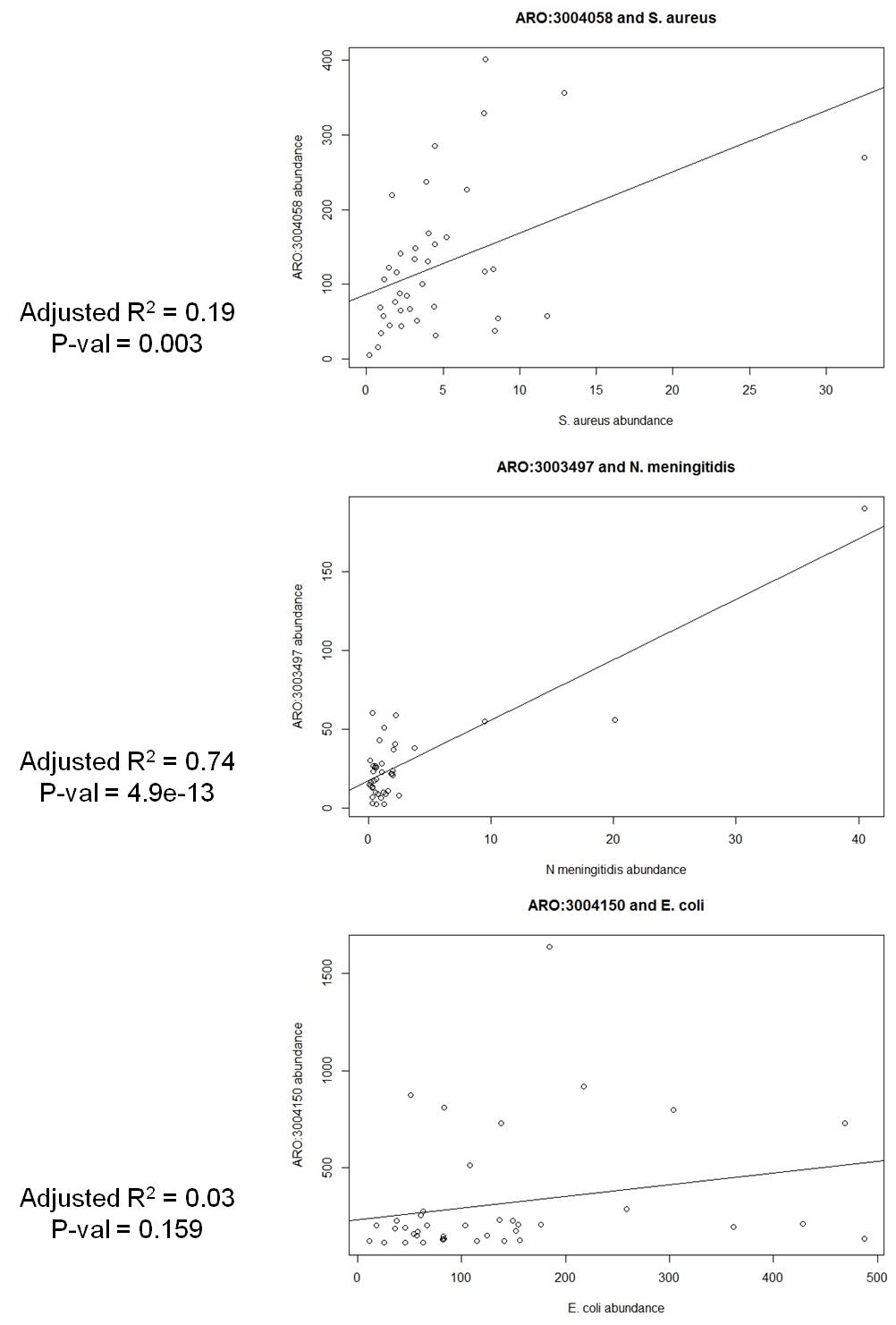
